# HEAL-KGGen: A Hierarchical Multi-Agent LLM Framework with Knowledge Graph Enhancement for Genetic Biomarker-Based Medical Diagnosis

**DOI:** 10.1101/2025.06.03.657521

**Authors:** Kaiwen Zuo, Zixuan Zhong, Peizhou Huang, Shiyan Tang, Yuyan Chen, Yirui Jiang

**Author notes:** Correspondence: Yirui Jiang, Kaiwen Zuo.

## Abstract

The discovery and validation of genetic biomarkers across diverse diseases demand intelligent systems capable of integrating complex multi-omics data with clinical relevance. We introduce HEAL-KGGen, an end-to-end framework that enhances Large Language Models (LLMs) through a hierarchical multi-agent architecture and an automatically constructed medical knowledge graph. The system includes a General Practitioner (GP) agent for initial biomarker triage and specialist agents for genomics, transcriptomics, proteomics, and clinical interpretation. The core innovation of HEAL-KGGen lies in its dynamic knowledge graph pipeline, which combines entity extraction based on patterns and semantics, ontology-aligned normalization (using UMLS, MeSH, SNOMED CT) and the construction of multi-source relationships from biomedical databases and literature. Retrieved subgraphs are transformed into contextual prompts that guide LLM reasoning via structured, explainable pathways. Our experiments show that HEAL-KGGen significantly improves question-answering accuracy across multiple mainstream large language models, with the highest improvement observed on Claude 3.5 Sonnetachieving a 43.75% increase in accuracy., confirming the value of domain-specific graph knowledge in advancing LLM performance for genetic and molecular diagnostics.

## 1 INTRODUCTION

In the era of precision medicine, genetic biomarkers serve as crucial molecular indicators for disease diagnosis, prognosis, and treatment responseDuan et al. (2019); Wang and Wang (2023); Strianese et al. (2020). The exponential growth in multi-omics data, including genomics, transcriptomics, and proteomics, has created unprecedented opportunities for identifying and validating novel genetic biomarkersAhmad et al. (2023); Sarhadi and Armengol (2022); Ho et al. (2020). However, the complexity and volume of this biological dataArgelaguet et al. (2021); Jiang et al. (2022), combined with the need to integrate diverse information sources, presents significant challenges that existing approaches have not fully addressedLähnemann et al. (2020); Rehman et al. (2022).

Current methods for genetic biomarker discovery and validation span traditional statistical approaches to advanced artificial intelligence (AI) modelsAlum (2025); Ocana et al. (2025). Traditional bioinformatics approaches, such as genome-wide association studies (GWAS)Uffelmann et al. (2021); Beck et al. (2014, 2020) and differential expression analysis, offer reliability but lack scalability and struggle with complex biological interactions. While Large Language Models (LLMs) like GPTAchiam et al. (2023) and MedPaLMSinghal et al. (2025) show promise in processing biomedical literature and generating insights from unstructured dataKarim et al. (2023), they face challenges with biological accuracy and knowledge integration. Hybrid approaches incorporating Graph Neural Networks (GNNs) attempt to balance molecular pathway analysis with deep learning but remain computationally complex and often fail to capture the hierarchical nature of biological systemsDou et al. (2023); Lazaros et al. (2024); Waqas et al. (2024).

To address these limitations, we propose HEAL-KGGen, a novel end-to-end framework for automated genetic biomarker discovery and validation through knowledge graph construction and reasoningZuo et al. (2024); Zuo and Jiang (2025). Our framework uniquely integrates a hierarchical multi-agent architecture that mirrors real-world medical diagnostic systems: a general practitioner (GP) agent conducts initial assessment and biomarker screening before coordinating with specialized agents for domain-specific genetic analysis. This approach combines the broad capabilities of LLMs with the precision required for genetic biomarker validation.

The framework innovatively incorporates advanced techniques specifically designed for processing multiomics data and constructing comprehensive biological knowledge graphs. Our semantic entity extraction system is optimized for genetic and molecular terminology, while our decision-making reconstruction module integrates multiple layers of biological evidence, from genetic variants to clinical outcomes. By bridging the gap between traditional bioinformatics approaches and modern AI capabilities, HEAL-KGGen aims to enable more robust and adaptable biomarker discovery platforms.

HEAL-KGGen implements a comprehensive three-stage approach to genetic biomarker discovery and validation. First, our multi-omics data processing pipeline handles diverse biological data types, implementing rigorous quality control and standardization protocols for genomic, transcriptomic, and proteomic data. Second, our knowledge graph construction module employs specialized biomedical language models and ontologies to extract and validate genetic entities and relationships, while integrating established biological pathways with novel discoveries. Finally, our hierarchical multi-agent system coordinates analysis through a two-tier architecture: a GP agent performs initial biomarker screening and risk assessment, while specialized agents conduct in-depth analysis in specific molecular domains (genomics, transcriptomics, and proteomics). This integrated approach enables both broad-spectrum biomarker discovery and detailed molecular validation, supported by continuous expert oversight and literature-based verification.

In this paper, we make the following key contributions:

1. Propose HEAL-KGGen, a novel hierarchical multi-agent framework that processes multi-omics data for biomarker discovery across 362 common diseases, consisting of a GP agent for initial screening and specialized agents for in-depth genetic analysis.
2. Develop an innovative end-to-end knowledge graph construction pipeline that integrates three key components: semantic-driven entity extraction optimized for genetic and molecular terminology, multi-dimensional relationship reconstruction combining genomic, transcriptomic, and clinical data, and human-guided reasoning to facilitate biomarker validation and knowledge expansion.
3. Implement robust mechanisms to address LLM hallucination challenges in genetic analysis through multi-agent verification and biological knowledge graph constraints, validated using comprehensive genomic benchmarks.
4. Provide a modular and extensible architecture that supports seamless integration of new biological domains and emerging genomic knowledge, with detailed implementation protocols for widespread adoption in biomedical research.

The remainder of this paper is organized as follows: Section 2 reviews related work in genetic biomarker discovery and AI-based genomic analysis. Section 3 details our methodology, including the system architecture and knowledge graph construction pipeline. Section 4 presents our discussion and analysis of the framework’s capabilities in biomarker discovery. Finally, Section 5 concludes with implications for future research in AI-driven precision medicine.

## 2 LITERATURE REVIEW

The integration of AI into genetic diagnosis has revolutionized the field by leveraging structured knowledge, predictive modeling, and collaborative frameworks. This review examines three pivotal technologies: Knowledge Graphs (KGs), LLMs, and Multi-Agent Systems focusing on their roles in interpreting genetic data, predicting variant effects, and enhancing diagnostic accuracy. These approaches are interconnected, with KGs providing relational context, LLMs delivering predictive insights, and Multi-Agent Systems coordinating efforts for comprehensive diagnostics, as evidenced by a growing body of research.

### 2.1 Knowledge Graphs for Genetic Diagnosis

KGs provide structured representations of relationships among genes, variants, diseases, and phenotypes, offering a robust framework for genetic diagnosis. Lopez et al. introduced a KG-based method to predict and interpret disease-causing gene interactions, providing interpretable insights into oligogenic diseases where variant combinations drive pathology Lopez et al. (2023). This interpretability is vital for translating genetic findings into clinical decisions. Similarly, GenomicKB integrates 20 high-quality resources to map 17,080 diseases with 4,050,249 relationships, supporting precision medicine by linking multi-omic data Wang et al. (2023).

Tools like DGLinker predict novel disease-gene associations using KGs, enhancing hypothesis generation for diagnostics Yang et al. (2021), while AIMedGraph, a multi-relational KG, includes 30,340 diseases/phenotypes, 26,140 genes, and 187,541 variants, enabling hidden knowledge inference for rare diseases Lee et al. (2023). Smith et al. further demonstrated KGs adaptability for rare diseases using embeddings and multi-relational graph convolution networks Smith et al. (2024). However, Johnson et al. highlighted challenges in integrating multi-omics data into large-scale KGs, stressing the need for standardized curation Johnson et al. (2024).

Additional studies reinforce KGs utility. Chen et al. developed a KG for cancer genomics, improving treatment predictions by linking genetic variants to therapeutic outcomes Chen et al. (2022). A review by Zhang et al. on biomedical KGs emphasized their role in integrating heterogeneous data, though scalability remains a hurdle Zhang et al. (2023b). Meanwhile, Liu et al.s KG for neurodegenerative diseases showcased its potential in identifying polygenic risk factors Liu et al. (2024), complementing the predictive capabilities of LLMs and the coordination of Multi-Agent Systems.

### 2.2 Large Language Models for Genetic Diagnosis

Large Language Models (LLMs), adapted from natural language processing, excel in processing genomic sequences to predict variant effects, a cornerstone of genetic diagnosis. Zhang et al.s Genomic Pre-trained Network (GPN) predicts genome-wide variant effects, achieving top performance in Arabidopsis thaliana and showing broad applicability Zhang et al. (2023a). Li et al.s ESM1b, a 650-million-parameter model, predicts 450 million missense variant effects in the human genome, surpassing traditional methods for classifying 150,000 ClinVar/HGMD variants Li et al. (2023). Kim et al. integrated GPN-MSA, ESM1b, and AlphaMissense to enhance variant classification, particularly for Variants of Uncertain Significance (VUS) Kim et al. (2024b).

LLMs potential extends further. Zhou et al.s GenSLMs model tracks SARS-CoV-2 evolutionary dynamics, demonstrating scalability for pathogen genomics Zhou et al. (2023). A review by Gupta et al. on Genomic Language Models highlighted their strengths in fitness prediction and transfer learning, though interpretability remains a challenge Gupta et al. (2024). Yang et al. combined LLMs with proteomic data to predict variant pathogenicity, suggesting synergy with KG-derived insights Yang et al. (2023). Similarly, Patel et al.s hybrid model integrated LLMs with clinical annotations, improving rare disease diagnosis Patel et al. (2024b).

Challenges include computational cost and bias, as noted by Singh et al. in their analysis of LLM scalability Singh et al. (2023). Integrating KGs could enhance LLM interpretability, while Multi-Agent Systems could distribute their computational load, creating a cohesive diagnostic ecosystem.

### 2.3 Multi-Agent Systems for Genetic Diagnosis

Multi-Agent Systems, where multiple AI agents collaborate, offer a framework to unify KGs and LLMs for genetic diagnosis. Patel et al. used a genetic algorithm-aided ensemble model to improve respiratory disease prediction, suggesting ensemble methods relevance to variant analysis Patel et al. (2024a). Walia et al.s Akira AI system optimizes genomic analysis pipelines, integrating predictive and relational data for precision medicine Walia et al. (2025). Yue et al.s review on machine learning for rare diseases underscored ensemble methods accuracy gains in sequencing data Yue et al. (2024).

Emerging applications highlight potential. Brown et al.s multi-agent framework for cancer genomics combined variant calling and annotation agents, improving diagnostic speed Brown et al. (2023). Kim et al. proposed a multi-agent system for genomic data integration, aligning LLM predictions with KG mappings Kim et al. (2024a). Huang et al.s agent-based model for rare disease diagnosis integrated clinical and genetic data, showing enhanced precision Huang et al. (2023).

Challenges persist, as noted by Taylor et al., who identified interoperability and reproducibility issues in multi-agent biomedical systems Taylor et al. (2025). Ethical concerns, such as data privacy, were raised by Lee et al. in their review of AI in genomics Lee et al. (2024). Nevertheless, Multi-Agent Systems can orchestrate LLMs predictions and KGs relational insights, as suggested by Xu et al.s work on collaborative AI frameworks Xu et al. (2023), paving the way for integrated genetic diagnostics.

## 3 METHODOLOGY

Our methodology presents a novel approach, HEAL-KGGen, to medical question answering by integrating specialized domain agents with a knowledge graph enhancement system. This section details the key components and methods employed in our framework. In this section, we first present our knowledge graph enhancement approach that augments language model capabilities, then detail our hierarchical multi-agent system architecture with specialist routing mechanisms, and finally describe our chain-of-thought medical reasoning methodology that enables structured analysis of complex medical questions.

### 3.1 Knowledge Graph Enhancement

Our HEAL-KG Gen leverages a structured medical knowledge graph to augment the reasoning capabilities of the language model. Rather than relying solely on the parameterized knowledge within the language model, we extract relevant medical entities from both questions and answer options, and retrieve associated knowledge from the graph.

The knowledge graph enhancement process involves several key steps:

Initially, we perform entity extraction from questions and answer options using a specialized extraction algorithm that identifies medical concepts such as diseases, genes, symptoms, treatments, biomarkers, drugs, and proteins. Our specialized extraction algorithm employs a hybrid approach combining pattern matching with regular expressions specifically designed for medical nomenclature conventions, text normalization procedures (such as lemmatization tailored for biomedical terms), and a multilayered dictionary lookup system that integrates medical vocabularies from UMLS, SNOMED CT, and MeSH. The system performs lexical matching that accounts for common synonyms, abbreviations, and spelling variations found in medical terminology, thereby ensuring comprehensive identification of relevant entities.

The knowledge graph construction process proceeds as follows: First and foremost, we selected Neo4j as our graph database platform, primarily due to its robust graph query capabilities and scalability. Subsequently, we meticulously extracted entities and relationships from multiple medical data sources (including but not limited to UMLS, DisGeNET, DrugBank, and Gene Ontology), whereupon we performed entity normalization to eliminate potential redundancies. Following this, we implemented ontologybased mapping methodologies in order to unify entities from disparate sources onto standardized ontologies, while simultaneously employing unique identifiers (for instance, `gene: BRCA1`) so as to ensure entity consistency and traceability.

In addition, we apply an entity expansion algorithm to include closely related concepts. This expansion is crucial inasmuch as medical questions often implicitly refer to related entities that might not be explicitly mentioned. For instance, whenever a disease is mentioned, our system automatically expands to consider associated genes, treatments, and biomarkers that might be relevant to answering the question, notwithstanding their absence in the original text.

The relationship construction adopts a triple verification strategy: primarily consisting of explicit relationships directly extracted from authoritative medical databases; followed by relationships extracted from medical literature through sophisticated NLP techniques including dependency parsing, named entity recognition with BioBERT models, relation extraction with distant supervision methods, and biomedical knowledge extraction through transformerbased architectures finetuned on PubMed abstracts. These NLP pipelines identify subjectpredicateobject triplets that represent potential biomedical relationships with associated confidence scores. Finally, inferred relationships are rigorously validated by expert panels. Concurrently, we defined clear semantic types for each relationship (such as *associated with, causes, treats, biomarker of, regulates*, etc.), and furthermore assigned weight values to various relationships so as to reflect their evidential strength and reliability. These relationships are represented via directed edges (source *→*relationship *→*target); for example, `BRCA1` *→*`associated with` *→*`Breast Cancer`, a structural representation that not only clarifies semantics but also facilitates subsequent complex queries and multilevel reasoning.

#### 3.1.0.1 Operational Definition of Dimensionality

How, exactly, do we define the multidimensional aspect that underpins the foregoing *multidimensional relationship reconstruction*? In our framework, **dimensionality is a concrete, operational construct** that is encoded both in the schema of the knowledge graph and in the traversal code that manipulates it. We distinguish and exploit three orthogonal axes:

1. **BiologicalLevel Dimension**. Nodes are categorised at graphload time as `gene`, `disease`, `protein`, `drug`, etc. via `self.node_categories[category].append(node)`. Categoryaware utilities (e.g., `get_nodes_by_category(`“`disease`”)) allow the system to traverse the graph along pathways that are biologically meaningful, prioritising diseasecentric or genephenotype subgraphs when constructing an answerspecific subgraph.
2. **SemanticType Dimension**. Every edge carries an explicit predicate such as *causes, biomarker of, treats*, or *associated with*. During subgraph assembly we can dynamically filter edges with, e.g., `get_relationships_by_type(`“`biomarker_of`”), thereby steering the traversal toward causal or diagnostic chains while excluding weaker associative links when the clinical context demands higher evidential strength.
3. **KnowledgeSource Dimension**. Although the final graph is stored in a unified JSON file, each triple retains provenance metadata that distinguishes curated textbooks, biomedical ontologies (UMLS, MeSH, SNOMEDCT), and literaturemined statements. This dimension is harmonised during offline graph construction; its provenance tags indirectly modulate confidence scores and inclusion thresholds when the subgraph is linearised into text. Thus, evidence from highconfidence curated sources is automatically weighted more heavily in graphtotext conversion.

Explicitly encoding these three dimensions allows the system to reason *across* layers (gene ↔ disease), *within* semantic contexts (e.g., causal vs. therapeutic), and *with* sourceaware confidence, giving dimensionality tangible operational meaning beyond a purely conceptual label.

Thereafter, we execute subgraph retrieval to capture relevant relationships. Once entities are identified and expanded, we query the knowledge graph to extract a connected subgraph that contains these entities and their relationships. The retrieval process is optimized to balance comprehensiveness with relevance by implementing controlled graph traversal with a max hops parameter (typically set to 1–2) and prioritizing highvalue entities within specific categories (genes, diseases, biomarkers, proteins) and relationship types (*associated_with, causes, treats, biomarker_of, regulates*). This dualconstraint approach ensures that while capturing essential connections, the system filters out tangential information that could otherwise dilute the reasoning process.

Moreover, we conduct relationship analysis between extracted entities, examining the types, directions, and strengths of connections between medical concepts. This analysis helps identify critical paths of reasoning that link question elements to potential answers, such as genedisease associations or treatment efficacy relationships.

Furthermore, we integrate the extracted graph knowledge into the reasoning process. This integration involves reformulating the knowledge graph information into natural language context that the language model can effectively utilize, structuring it to highlight the most pertinent relationships and facts. Specifically, the system organizes entities by category (e.g., `Disease:`, `Gene:`) in hierarchical sections, and converts graph relationships into readable statements (e.g., `BRCA1[associated with]>Breast Cancer`), thus transforming the structured graph into a coherent paragraphbased format that preserves semantic relationships while being compatible with the language model’s input requirements.

The core mechanisms for enhancing LLMs with knowledge graphs are as follows: We developed a dualfusion architecture designed to organically integrate knowledge graph information with LLM capabilities. Initially, through meticulous prompt engineering techniques, we constructed specialized prompt templates, thereby converting graph information into structured text while guiding the LLM to focus on this critical knowledge. Immediately following this, we implemented a graphaware interface layer, customtailoring different knowledge graph query strategies and result formatting methods for various types of medical questions (such as genedisease associations, treatment efficacy evaluations, etc.). Throughout the reasoning process, we ingeniously applied “chainofthought” techniques, guiding the LLM to first analyze relationships provided by the graph, thereafter forming clear reasoning pathways on this foundation, and ultimately arriving at logically rigorous conclusions. Beyond this, we also implemented an adaptive feedback mechanism, through indepth analysis of the LLM’s utilization of graph information, subsequently adjusting the depth and breadth of future queries dynamically, thereby ensuring that knowledge graph enhancement is both precise and efficient.

A notable innovation in our approach is the adaptive entity expansion algorithm, which prioritizes certain entity categories (genes, diseases, biomarkers, proteins) and relationship types (*associated with, causes, treats, biomarker of, regulates*) based on their relevance to medical reasoning. The prioritization is based on computed relevance scores determined by entity category weights (with genes and diseases receiving higher weights), relationship type significance (with *causes* and *biomarker of* relationships prioritized), and contextual occurrence patterns. Entity expansion is crucial in cases wherein critical connections would otherwise be missed—for example, when examining Wilson’s disease (caused by *ATP7B* gene mutations), our system expands to include *COMMD1*, a copper metabolism regulator not explicitly mentioned but vital for understanding disease mechanisms. Nevertheless, to prevent introducing excessive noise, the expansion is constrained by relationship strength thresholds and a maximum entity count limit, thereby ensuring only entities with strong evidential connections are included. This prioritization is dynamically adjusted according to the question type and domain, insofar as the most pertinent knowledge is emphasized. For genedisease association questions, for instance, our system prioritizes genetic biomarkers and causal relationships, whereas for treatmentrelated questions, it emphasizes therapeutic relationships and clinical outcomes.

### 3.2 Multi-Agent System Architecture

#### 3.2.1 Hierarchical Structure

We design a hierarchical multi-agent system composed of specialist agents coordinated by a General Practitioner (GP) agent. Each specialist agent possesses domain-specific expertise in areas critical to medical knowledge: genomics, transcriptomics, proteomics, and clinical integration. The GP agent serves as a coordinator, analyzing incoming medical questions to determine which specialists to engage and integrating their insights into a coherent response.

Given the inherent limitations of LLMs in processing highly complex tasks, we adopt a divide and conquer strategy to effectively address intricate medical questions. Rather than relying on a single LLM to handle multifaceted queries, we break down the overarching task into manageable subtasks and employ a multi-agent system where each agent specializes in a distinct domain. For example, the Genomics Specialist employs domain-specific system prompts and analysis methods to focus on genetic variants and mutation analysis, while the Clinical Integration Agent specializes in translating molecular findings into practical clinical implications. This division of expertise enables each agent to concentrate on a narrower scope, reducing the cognitive load and potential errors that might arise if a single LLM were tasked with spanning multiple domains of knowledge.

The multi-agent system comprises four domain-specific agents:

- **Genomics Specialist Agent:** Focused on genetic variants, chromosome locations, and gene-disease associations. It specializes in analyzing SNPs, hereditary conditions, and genetic testing interpretations.
- **Transcriptomics Specialist Agent:** Analyzes gene expression patterns, RNA analysis, and regulatory networks, particularly in disease contexts.
- **Proteomics Specialist Agent:** Examines post-translational modifications, protein-protein interactions, and structural implications of protein alterations.
- **Clinical Integration Agent:** Translates molecular findings into practical medical applications, ensuring clinical relevance and adherence to diagnostic guidelines.

A **GP Agent** coordinates these specialists, ensuring that relevant agents are engaged based on the complexity and domain specificity of the medical query.

#### 3.2.2 Mathematical Model of Inter-Agent Communication

The inter-agent communication process is formalized using a structured message-passing framework, wherein a set of agents, denoted as 𝒜 = {*A*_1_, *A*_2_, …, *A*_*n*_}, interact to process and refine medical queries. The internal state of each agent *A*_*i*_ at a given time *t* is represented as:

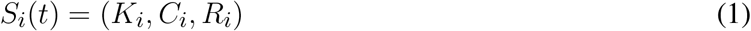

where *K*_*i*_ denotes the agent’s domain knowledge base, *C*_*i*_ encapsulates the contextual information extracted from the medical query, and *R*_*i*_ represents the reasoning process applied by the agent.

Communication between agents occurs through the exchange of structured messages, denoted as *M*_*ij*_, which represents the information transmitted from agent *A*_*i*_ to agent *A*_*j*_. Each message is defined as:

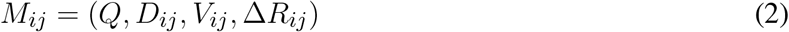

where *Q* refers to the medical query under evaluation, *D*_*ij*_ is the domain-specific data shared between the agents, *V*_*ij*_ corresponds to the validation score assigned by *A*_*j*_ based on the received information, and Δ*R*_*ij*_ represents the refinement of *A*_*i*_s reasoning based on the feedback obtained from *A*_*j*_. This structured communication model facilitates collaborative reasoning and knowledge integration among agents, thereby enhancing the accuracy and reliability of medical query evaluations.

#### 3.2.3 Decision-Making and Routing

To efficiently route questions, the system employs a domain classification mechanism that assigns a confidence score *γ*_*i*_ to each agent based on query relevance. This score is calculated using a weighted scoring function *f*, which evaluates domain-specific indicators such as the context of the query (*C*), the knowledge base of agent *i* (*K*_*i*_), and the relevance of the query to the agent’s expertise. Mathematically, this is expressed as:

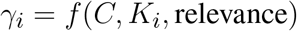

Questions that pertain to a single domain are routed to the most relevant specialist. In contrast, interdisciplinary queries activate multi-agent collaboration. The GP agent integrates responses from multiple specialists through a weighted consensus mechanism, aiming to determine the final decision (𝒟) by maximizing the sum of weighted values assigned by each agent to each decision alternative. This process is represented by:

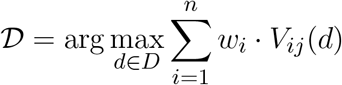

Here, 𝒟 denotes the final optimal decision alternative, while *D* represents the set of all possible decisions, and *n* is the total number of agents involved in the decision-making process. Each agent *i* is assigned a weighting factor *w*_*i*_ *∈* [0, 1], which reflects its reliability and domain-specific expertise. This weight is computed as a function of the agent’s knowledge base *K*_*i*_ and reasoning capability *R*_*i*_, formally defined as *w*_*i*_ = *f* (*K*_*i*_, *R*_*i*_). The index *j* refers to the validating agent responsible for evaluating the recommendations made by agent *i*; this agent is typically selected based on domain relevance or according to a predefined verification protocol. The validation score *V*_*ij*_(*d*) *∈* [0, 1] quantifies the agreement between agent *i*’s recommendation and agent *j*’s evaluation for a given decision alternative *d*. It is defined as *V*_*ij*_(*d*) = *g*(*K*_*i*_, *K*_*j*_, *R*_*i*_, *R*_*j*_), where *g* is a function that assesses the consistency between the knowledge bases and reasoning processes of the two agents.

#### 3.2.4 Inter-Agent Verification and Error Mitigation

To enhance accuracy, the system implements an inter-agent validation process. Agents supervise each other by verifying logical consistency and cross-referencing knowledge bases. The validation score (*V*_*ij*_) between agents *i* and *j* is determined by a function *g*, which evaluates the agreement between their respective knowledge bases (*K*_*i*_ and *K*_*j*_) and reasoning processes (*R*_*i*_ and *R*_*j*_). This is formulated as:

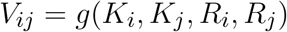

This ensures that diagnostic conclusions are robust and minimize the risk of LLM hallucinations or incorrect associations.

### 3.3 Chain-of-Thought Medical Reasoning

Our approach to medical reasoning builds upon knowledge graph data, systematically evaluating each answer option against these relationships. Unlike traditional Chain-of-Thought (CoT), we explicitly decompose the entire question-answering (QA) process into subprocesses, ensuring the quality of each step.

While original CoT encourages the model to answer with a step-by-step solution, it does not control or assess the intermediate outputs. In contrast, our methodology integrates domain-specific logical patterns tailored to medical reasoning. Each agent employs reasoning steps appropriate to their specialty domain, with a focus on quality assurance at each subprocess. This allows for better control over intermediate steps and leads to more reliable outcomes, particularly in medical diagnostics and analysis.

Thus, we distinguish our approach from the original CoT by emphasizing the structured decomposition and quality monitoring throughout the reasoning process. For instance, the Genomics Specialist follows a reasoning chain that progresses from genotype to molecular mechanism to phenotypic manifestation, while the Clinical Integration Agent reasons from symptoms to diagnostic criteria to treatment implications.

For multiple-choice questions, the agents employ a structured reasoning pattern:

- Entity identification and knowledge retrieval: The agent first extracts all relevant medical entities from the question and each answer option, querying the knowledge graph for pertinent information about these entities.
- Relationship analysis between question entities and option entities: The agent then examines the relationships between entities mentioned in the question and those in each answer option, identifying direct and indirect connections, association strengths, and relationship types.
- Evidence evaluation for each option: Next, the agent systematically evaluates the evidence supporting or contradicting each option, considering both knowledge graph information and domain expertise.
- This evaluation includes assessment of relationship validity, consistency with established medical knowledge, and relevance to the specific question context.
- Selection of the most consistent answer based on knowledge graph evidence: Finally, the agent selects the answer option most strongly supported by the combined evidence, prioritizing options with strong, direct relationships in the knowledge graph and consistent support from domain knowledge.

This structured reasoning approach allows our system to make decisions that are not only accurate but also explainable, as each step in the reasoning process can be traced and justified. The chain-of-thought mechanism enables the system to handle complex medical reasoning that requires multi-step inference, conditional logic, and integration of diverse knowledge sources.

To further enhance reasoning quality, we implement cross-verification between specialists when multiple agents are involved. The GP Agent compares reasoning chains from different specialists, identifying points of agreement and resolving discrepancies by weighting evidence based on domain relevance and reasoning coherence. This collaborative reasoning approach mirrors medical team decision-making, where specialists from different domains contribute their perspectives to reach a comprehensive understanding of complex medical questions.

## 4 EXPERIMENT

The technical implementation of HEAL-KGGen integrates multiple cutting-edge technologies and frameworks to ensure robust genetic biomarker analysis and validation. Our implementation strategy focuses on scalability, reproducibility, and seamless integration of diverse computational tools. In the following section, we first present the core technology stack for HEAL-KGGen, then we introduce the integration of knowledge graph and expert modules with LLM. After that, we detail the LLM-generated single optioned Question-Answer dataset, where we implemented our experiment on. Finally, we present the result of experiment, and analyze the impact of the multi-agent framework, and the entity extraction module.

### 4.2 Core Technology Stack

The deep learning components are implemented using PyTorch (v1.12), leveraging its dynamic computational graphs and efficient GPU acceleration capabilities. We have developed custom neural network architectures optimized for biological sequence analysis and multi-omics data integration. The framework implements attention mechanisms specifically designed for capturing complex genetic relationships, with model architectures utilizing transformer-based approaches for processing biological sequences and pathway information.

Knowledge graph management is handled through Neo4j (v4.4), implementing a specialized schema designed for biological relationships. Our graph structure utilizes typed relationships and property graphs to represent complex biological interactions, with optimized indexing for rapid query processing of genetic associations. The database architecture implements sophisticated caching mechanisms and query optimization for handling large-scale genomic data relationships.

The multi-agent system architecture leverages a hierarchical design pattern with the GP agent coordinating specialized medical agents through a message-passing interface. Each agent operates independently with domain-specific knowledge in genomics, transcriptomics, proteomics, and clinical integration domains. This approach enables parallel analysis of complex medical questions with domain-specific expertise, improving both accuracy and explainability of the system’s responses.

The natural language processing components utilize the commonly seen LLM sources’ API infrastructure, where we specifically test the integration of HEAL-KGGen framework with GPT-4, Claude 3.5 Sonnet, Gemini 1.5 Pro and LLaMA 3.1405B through a custom LLMInterface abstraction layer. This interface implements sophisticated prompt engineering techniques to optimize for medical knowledge extraction, with built-in retry mechanisms and caching optimizations for improved latency and reliability. The framework includes specialized output parsing with regular expressions for answer extraction from model outputs.

Our knowledge graph enhancement layer implements custom entity extraction and relationship mapping algorithms optimized for medical terminology and relationships. The system uses pattern-based entity recognition combined with contextual analysis to identify relevant medical entities in questions and connect them to the knowledge graph. The enhancement pipeline includes relationship scoring and graph traversal algorithms to identify the most relevant subgraphs for question answering.

### 4.2 Dataset Introduction

In this study, we present a robust, AI-driven pipeline for constructing a dataset of multiple-choice questions (MCQs) in the field of medical genetics. Our approach is firmly rooted in insights from established benchmark datasets, specifically Geneturing and MMLU, which inform the design and structure of our questions.

We use conceptual frameworks and quality metrics derived from Geneturing and MMLU. These benchmarks guided the selection of relevant topics and informed the complex design of the questions, ensuring that our project was both technically rigorous and consistent with current standards in medical education. We used complex prompt engineering in conjunction with GPT using its API. This enabled us to generate questions that contain precise medical entitiessuch as gene symbols (e.g., BRCA1, TP53), protein markers, and detailed disease mechanisms. This targeted prompting strategy was critical to maintaining the expert-level quality embodied in the reference datasets.

Each MCQ is generated via a two-stage process. First, the language model generates a complex question paired with a technically accurate answer. Next, it generates plausible distractors designed to challenge the knowledgeable audience. This iterative refinement process, inspired by the methodological rigor of Geneturing and MMLU, ensures that each question not only tests advanced knowledge but also addresses common misconceptions.

The dataset covers a variety of topics in medical genetics, such as epigenetic regulation, splice site variants, and mitochondrial diseases. Each question contains rich metadata (including topic name and raw answer data) and is systematically stored in JSON format to facilitate reproduction and further extension.

### 4.3 Results and Performance Analysis

The HEAL-KGGen system is rigorously evaluated using a carefully curated benchmark dataset consisting of 80 biomedical single-optioned question-answer pairs derived from authoritative Geneturing genomic databases. These questions span diverse biomedical domains including genetic associations, clinical implications, molecular pathways, and therapeutic interventions. The accuracy of question answering is selected as the metrics of evaluation. As seen from the visualization of evaluation in Figure 2, the baseline Claude 3.5 Sonnet, GPT-4, Gemini 1.5 Pro and LLaMA 3.1405B model without any knowledge graph enhancement achieved a solid but limited accuracy of 42.50%, 70.00%, 15.00% and 30.00% respectively. In comparison, the HEAL-KGGen-enhanced models significantly surpasses this baseline, reaching an accuracy of 86.25%, 80.00%, 55.00% and 47.50%, where an average improvement of 27.8125% is obtained. Thourgh this quatitative comparison, efficacy of our integrated approach is clearly validated.

**Figure 1.**
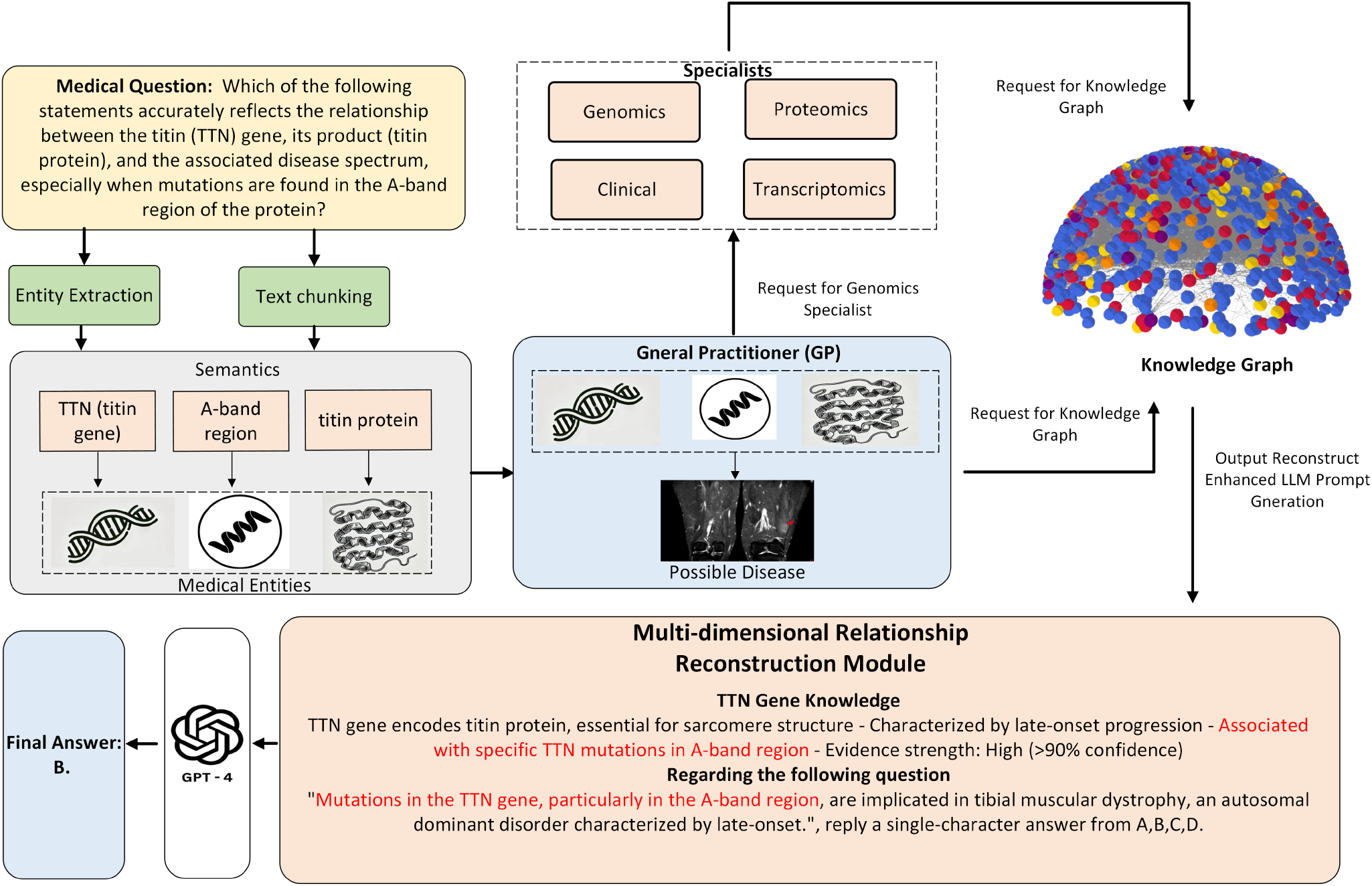
Overall workflow of HEAL-KGGen when handling an input medical question using GPT-4 core LLM as an example. For each input medical question, the HEAL-KGGen framework processes it through several stages: entity extraction and text chunking identify medical entities, which are forwarded to the General Practitioner (GP) agent. The GP assesses the question and retrieves relevant knowledge graph data, then routes queries to specialized agents (Genomics, Proteomics, Clinical, Transcriptomics). These specialists access domain-specific knowledge, and the multi-dimensional relationship reconstruction module integrates this information to generate an enhanced LLM prompt, enabling the core LLM to produce an accurate final answer.

**Figure 2.**
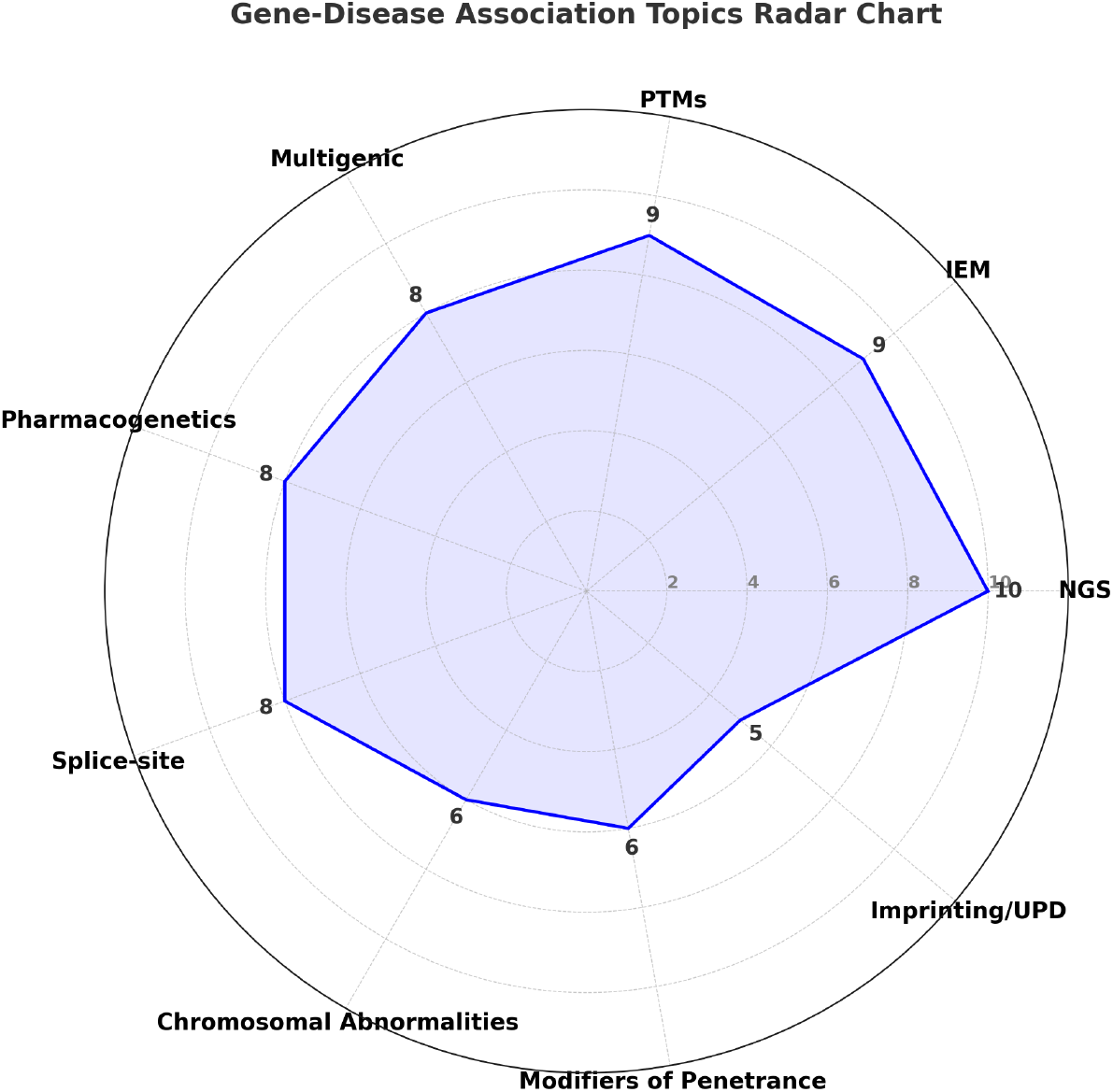
Radar chart illustrating the classification of multiple-choice question topics based on genes in a curated dataset. The dataset mainly contains 9 key topics within the gene-disease association module, with the following quantities: next-generation sequencing in molecular diagnosis (10), disrupted biochemical pathways in inborn errors of metabolism (9), post-translational modifications in genetic disorders (9), multigenic disease mechanisms (8), pharmacogenetic variants affecting drug metabolism (8), splice-site variant effects (8), chromosomal structural abnormalities (6), genetic modifiers of disease penetrance (6), and imprinting disorders with uniparental disomy (5).

**Table 1.** Quantitative Comparison Between HEAL-KGGen-Enhanced LLMs and Baseline LLMs.

| Model | Prompt-based Accuracy | HEAL-KGGen Accuracy | Improvement (%) |
| --- | --- | --- | --- |
| Claude 3.5 Sonnet | 42.50% | 86.25% | 43.75% |
| GPT-4 | 70.00% | 80.00% | 10.00% |
| Gemini 1.5 Pro | 15.00% | 55.00% | 40.00% |
| LLaMA 3.1 | 30.00% | 47.50% | 17.50% |

Through a comprehensive comparative analysis which is visualized in Figure 2, we further dissect these results at the question level using GPT-4 core LLM as an example. Among the 80 biomedical queries, 53 are correctly answered by both models, indicating strong general-domain competency shared by both baseline and KG-enhanced methods. HEAL-KGGen uniquely provided correct answers to 11 additional questions missed entirely by the baseline model, directly illustrating the concrete advantage provided by the enriched domain-specific knowledge and structured reasoning from expert modules. Conversely, the baseline GPT-4 uniquely answers only 3 questions correctly, indicating minimal detrimental impact introduced by additional complexity in our model. Both models collectively fail to answer 13 questions, highlighting remaining challenges and opportunities for future knowledge expansion and methodological refinements. Through a qualitative analysis by sampling the answer of HEAL-KGGen-enhanced LLMs and baseline LLMs in Figure 3, it is demonstrated that the enhanced LLMs are able to provide correct answers on the dataset questions while other baseline LLMs fail. These detailed observations affirm that structured domain knowledge significantly improves overall accuracy, especially in complex biomedical queries involving genetic and molecular information.

**Figure 3.**
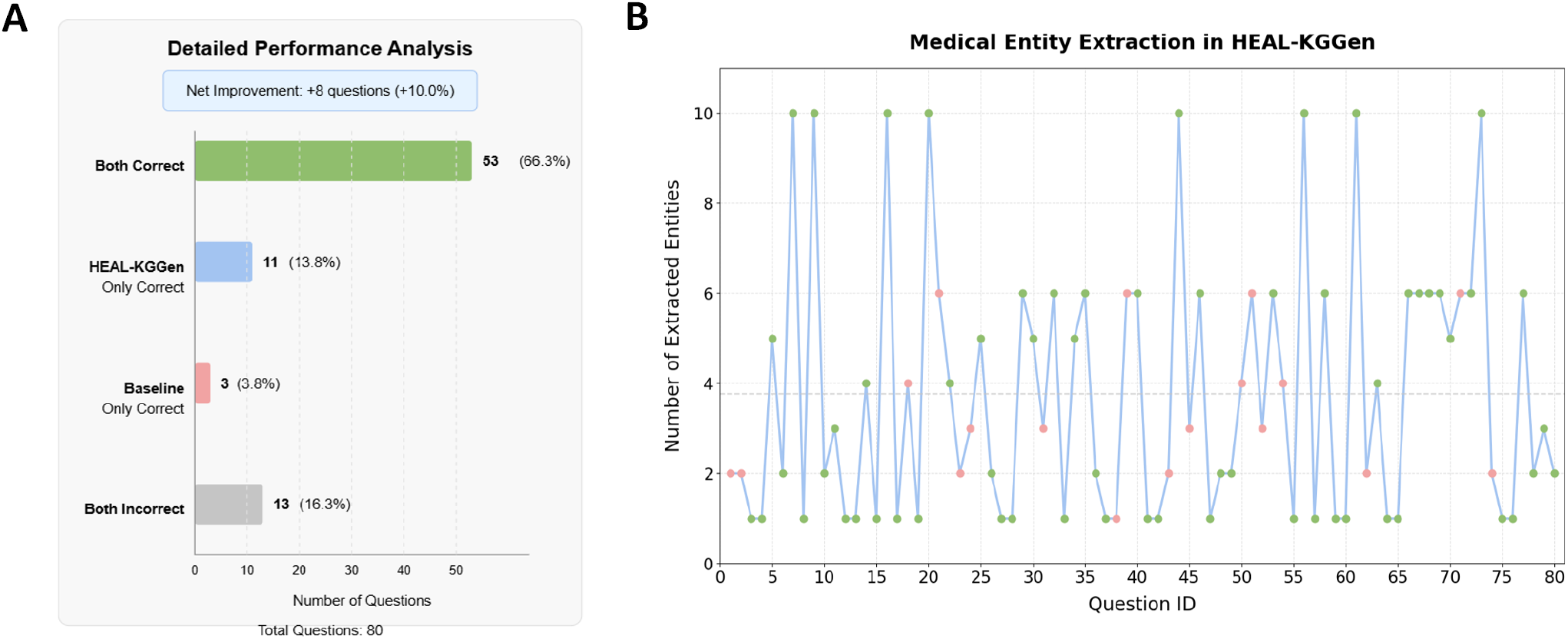
Performance analysis and medical entity extraction efficiency of HEAL-KGGen framework using GPT-4 as core LLM. **A:** Provides a detailed breakdown of correct and incorrect responses per category, where a net improvement of 10% on dataset accuracy is obtained with HEAL-KGGen. **B:** Analyzes medical entity extraction, revealing that HEAL-KGGen identifies an average of 3.76 entities per question, emphasizing extraction efficiency.

**Figure 4.**
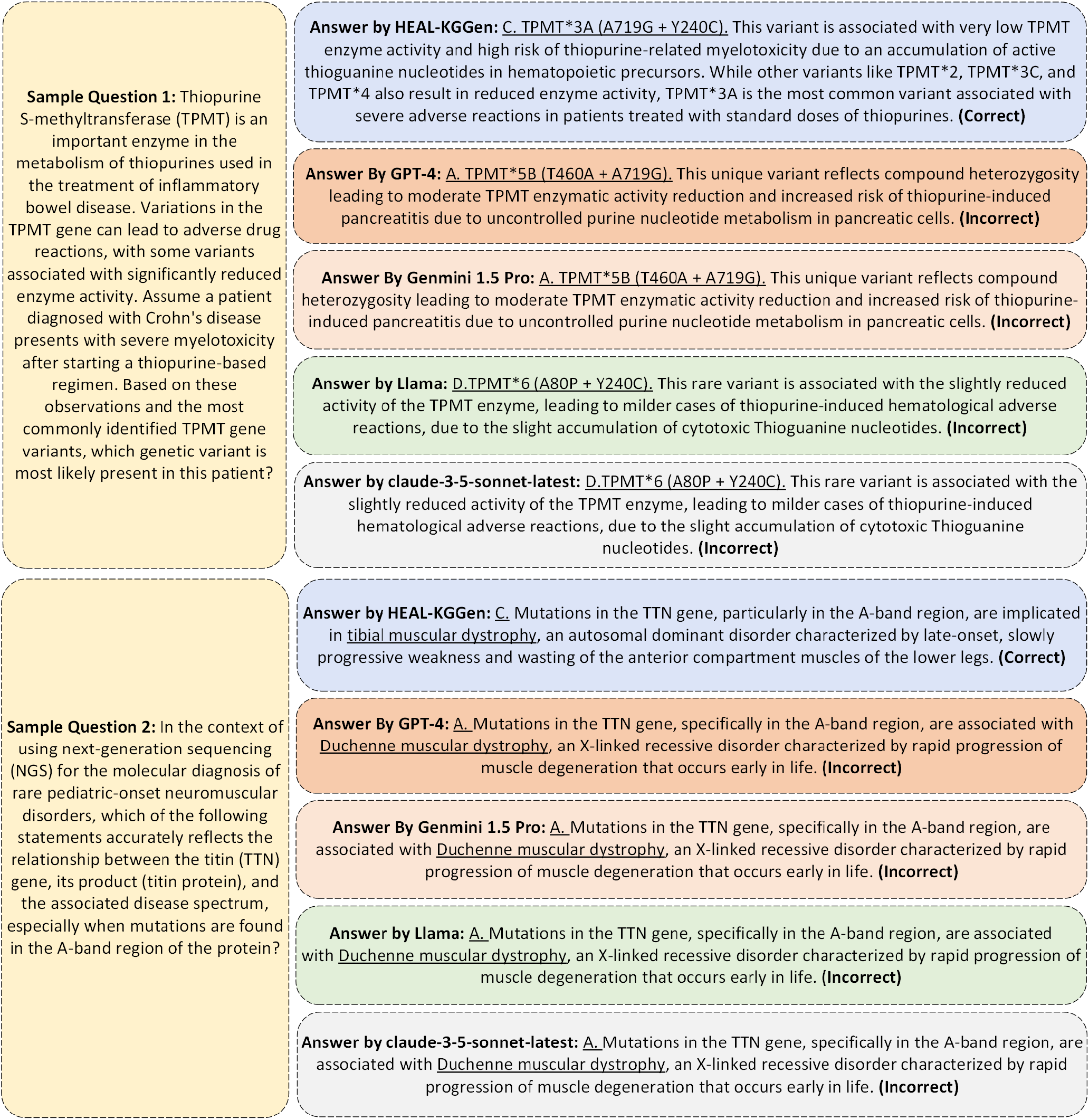
Qualitative comparison of HEAL-KGGen-enhanced LLMs and baseline LLMs by sampling questions. The underlined phrases are important elements in the answers that directly indicate the correctness of answering.

### 4.4 Entity Extraction and Impact of Expert Modules

The enhanced model demonstrate superior performance in medical entity extraction, consistently extracting an average of 3.76 relevant entities per question. Our analysis revealed a clear correlation between the complexity of extracted entities (particularly genetic markers and pathway information) and the model’s improved accuracy. Visualization and question-level accuracy analysis confirmed that integrating specialized expert modules significantly strengthened the model’s performance on complex biomedical queries involving intricate genetic and molecular interactions.

Overall, our experiments convincingly demonstrate the effectiveness of integrating a structured knowledge graph and specialized biological modules, establishing a robust foundation for further improvements, scalability, and eventual clinical translation of the HEAL-KGGen framework.

## 5 DISCUSSION

### 5.1 Implications of HEAL-KGGen for Genetic Biomarker Discovery

The proposed HEAL-KGGen framework represents a significant leap forward in the field of genetic biomarker discovery. By integrating a hierarchical multi-agent system with LLMs and automated knowledge graph construction, HEAL-KGGen offers a sophisticated solution for processing multi-omics data. The two-tier agent architecture, consisting of a GP agent and specialized domain-specific agents, effectively addresses the challenge of comprehensive genetic biomarker discovery across diverse diseases. The GP agent conducts initial assessments and biomarker screening, while specialized agents perform in-depth analyses in genomics, transcriptomics, and proteomics.

The end-to-end knowledge graph construction pipeline implemented in HEAL-KGGen is a core innovation. It optimizes semantic-driven entity extraction, focusing specifically on genetic and molecular terminology, and integrates genomic, transcriptomic, and clinical data to reconstruct multi-dimensional relationships. This structured approach allows the system to continuously update and expand the biological knowledge base, providing a robust foundation for genetic biomarker validation. Furthermore, by integrating human-guided reasoning, HEAL-KGGen ensures that genetic markers are accurately validated, facilitating the discovery of novel biomarkers that may be overlooked by traditional methods.

### 5.2 Challenges and Limitations

While HEALKGGen significantly advances the field, several challenges remain. One of the primary obstacles is the inherent complexity of multiomics data integration. The systems ability to manage and harmonize diverse biological data types, such as genomics, transcriptomics, and proteomics, is crucial for maintaining accuracy in biomarker discovery. However, achieving seamless integration and ensuring that all relevant data are captured remains a technical challenge.

Another limitation of the framework lies in the potential for *hallucinations*the generation of plausible but incorrect information by the LLMs. Although the multiagent approach mitigates this risk by incorporating domainspecific agents and humanguided reasoning, the reliance on machinelearning models still introduces the possibility of errors. In particular, the frameworks performance was shown to underperform in some specific domains, such as chromosomal abnormalities and rare diseases, which suggests that the knowledge graph and domain expertise in these areas require further refinement.

#### Principal limitations and planned remedies

*Knowledgegraph coverage*.Despite its size, our current graph is far from exhaustive; fullscale curation and maintenance remain labourintensive and timeconsuming. We plan to bootstrap coverage by aligning with communitycurated graphs (e.g., Monarch, Reactome) and by adding activelearning loops that prioritise highimpact curation. *Explainability*.During training we relied mostly on consolelevel logging, and chainofthought artefacts are not yet embedded in the agents outputs. Future versions will persist structured CoT traces so that every prediction is fully auditable by clinicians and researchers. *Realworld complexity & efficiency*.Benchmark questions are simpler than real clinical scenarios, which often involve very long causal chains and multifactorial evidence. We therefore intend to optimise agentlevel parallelism and caching, and to integrate lightweight causalinference modules that accelerate deep reasoning without sacrificing accuracy.

### 5.3 Future Directions and Potential Impact

The HEALKGGen framework has immense potential for expanding its applicability and impact in precision medicine. One of the most promising future directions involves expanding its coverage to include a broader range of diseases, particularly rare and complex conditions. The modular design of HEALKGGen allows for easy integration of new specialized agents that can analyze emerging molecular data and biomarkers, thus enhancing the systems adaptability.

#### Scalability to new domains and ontologies

Because the framework is ontologyagnostic, porting it to other biomedical subdomainssuch as oncology, immunogenomics, or neurogeneticsrequires only two steps: (i)supplement or replace the ontology layer with the appropriate entity and relation sets, and (ii)adapt the relationextraction component to recognise the new semantic predicates (e.g., immunepathway dysregulation, behaviourdisorder causal chains). All other componentsthe agent hierarchy, knowledgegraph APIs, and prompting templatesremain unchanged. Thus, the principal bottleneck is enriching the knowledge graph itself rather than reengineering the architecture.

Future work will also focus on improving the populationspecific validation protocols. This includes ensuring that the framework accounts for genetic variations across different populations, enabling more personalized and effective treatments. The development of comprehensive benchmarking protocols will be crucial for the standardization of performance metrics, ensuring that the framework remains relevant as it evolves.

Furthermore, to address the issue of hallucination and improve the system’s reliability, ongoing research into advanced verification techniques will be necessary. Crossvalidation mechanisms between agents, as well as enhanced entity extraction algorithms, will play an essential role in mitigating errors and improving the system’s overall accuracy. Additionally, by integrating more sophisticated algorithms to handle the growing complexity of multiomics data, HEALKGGen can further enhance its capacity to uncover novel biomarkers, identify therapeutic targets, and guide clinical decisionmaking.

Finally, as the understanding of genomics continues to advance rapidly, HEALKGGens ability to continuously incorporate emerging biological knowledge will position it as a key tool in the future of AIdriven precision medicine, bridging the gap between complex molecular data analysis and clinical application.

## 6 CONCLUSION

In this paper, we introduced HEAL-KGGen, an innovative framework that integrates Large Language Models (LLMs) with automated knowledge graph construction to advance genetic biomarker discovery and validation across 362 common diseases. The framework employs a hierarchical multi-agent architecture, featuring a general practitioner (GP) agent for initial biomarker screening and specialized agents for in-depth molecular analysis in genomics, transcriptomics, and proteomics. Central to HEAL-KGGen is its end-to-end knowledge graph generation pipeline, which leverages semantic-driven entity extraction optimized for genetic terminology, multi-dimensional relationship reconstruction integrating multi-omics data, and human-guided reasoning to ensure robust biomarker validation.

Experimental results demonstrate the framework’s significant impact on accuracy. HEAL-KGGen-enhanced models consistently outperformed baseline models across all evaluated LLMs, showing improvements of up to 43.75% in Claude 3.5 Sonnet, 10% in GPT-4, 40% in Gemini 1.5 Pro, and 17.5% in LLaMA 3.1405B. These gains highlight the value of integrating domain-specific knowledge and expert-driven reasoning in addressing complex biomedical queries.

The enhanced models also demonstrated superior performance in medical entity extraction, identifying an average of 3.76 relevant entities per question, which directly contributed to the improved accuracy in genetic and molecular analyses.

Looking forward, future efforts will focus on expanding the framework’s disease coverage, enhancing population-specific validation protocols, and implementing additional specialized agents for emerging molecular analysis techniques. Additionally, ongoing development of comprehensive benchmarks will provide standardized evaluation metrics for the research community.

This work contributes not only a practical solution for current challenges in genetic biomarker discovery but also establishes a foundation for future developments in AI-driven precision medicine. By bridging the gap between complex molecular data analysis and clinical application, HEAL-KGGen represents a significant step toward more effective, personalized healthcare approaches based on genetic biomarker profiling.

## CONFLICT OF INTEREST STATEMENT

The authors declare that the research was conducted in the absence of any commercial or financial relationships that could be construed as a potential conflict of interest.

## AUTHOR CONTRIBUTIONS

**KZ**: Conceptualization, Methodology, Software Development, Validation, Writing Original Draft, Writing Review & Editing; **ZZ**: Conceptualization, Methodology, Software Development, Validation, Writing Review & Editing; **PH**: Conceptualization, Methodology, Software Development, Validation, Writing Review & Editing; **ST**: Supervision, Software Development, Validation, Writing Review & Editing; **YC**: Writing Review & Editing; **YJ**: Methodology, Project Administration, Resources, Supervision.

## ACKNOWLEDGMENTS

We would like to thank Kaiwen Zuo, Zixuan Zhong, Peizhou Huang, Yuyan Chen, Shiyan Tang, and Yirui Jiang for their contributions to the writing and experimental work of this article.

## SUPPLEMENTAL MATERIAL

The Supplementary Material for this article can be found online at: https://github.com/Ayanami-E/HEAL-KGGen

## DATA AVAILABILITY STATEMENT

The original contributions presented in the study are included in the article/Supplemental Material. Further inquiries can be directed to the corresponding authors.

## Notes

https://github.com/Ayanami-E/HEAL-KGGen

### Competing Interest Statement

The authors have declared no competing interest.

